# Affective Touch Dimensions: From Sensitivity to Metacognition

**DOI:** 10.1101/669259

**Authors:** Mariana von Mohr, Louise P. Kirsch, Joey K. Loh, Aikaterini Fotopoulou

## Abstract

Touch can give rise to different sensations including sensory, emotional and social aspects. Tactile pleasure typically associated with caress-like skin stroking of slow velocities (1-10 cm/s) has been hypothesised to relate to an unmyelinated, slow-conducting C-tactile afferent system (CT system), developed to distinguish affective touch from the ‘‘noise’’ of other tactile information on hairy skin (the so-called ‘social touch hypothesis’). However, to date, there is no psychometric examination of the discriminative and metacognitive processes that contribute to accurate awareness of pleasant touch stimuli. Over two studies (total N= 194), we combined for the first time CT stimulation with signal detection theory and metacognitive measurements to assess the social touch hypothesis on the role of the CT system in affective touch discrimination. Participants’ ability to accurately discriminate pleasantness of tactile stimuli of different velocities, as well as their response bias, was assessed using a force-choice task (high versus low pleasantness response) on two different skin sites: forearm (CT-skin) and palm (non-CT skin). We also examined whether such detection accuracy was related to the confidence in their decision (metacognitive sensitivity). Consistently with the social touch hypothesis, we found higher sensitivity d’ on the forearm versus the palm, indicating that people are better at discriminating between stimuli of high and low tactile pleasantness on a skin site that contains CT afferents. Strikingly, we also found more negative response bias on the forearm versus the palm, indicating a tendency to experience all stimuli on CT-skin as ‘high-pleasant’, with such effects depending on order, likely to be explained by prior touch exposure. Finally, we found that people have greater confidence in their ability to discriminate between affective touch stimuli on CT innervated skin than on non-CT skin, possibly relating to the domain specificity of CT touch hence suggesting a domain-specific, metacognitive hypothesis that can be explored in future studies as an extension of the social touch hypothesis.

**Highlights:**

- Touch mediated by C-tactile (CT) afferents on hairy skin elicits pleasant sensations
- We combine for the first time CT stimulation with signal detection theory
- Better accuracy to detect pleasantness of tactile stimuli at CT optimal speeds on CT skin
- Higher confidence in ability to accurately distinguish affective touch on CT skin

## 1. Introduction

It is well known that tactile sensitivity varies across the body, which, among other things, depends on the neurophysiology of the skin (e.g., sensory afferent type and density). Human skin can be divided into two main categories, with different sensory afferent properties: glabrous skin (e.g., palm) and hairy skin (e.g., forearm). On the one hand, glabrous skin contains different types of myelinated mechanoreceptive afferents, which given their fast conduction velocity (60 m/s) provide the brain relatively fast sensory information about tactile stimulations, including their duration, texture, force, velocity and vibration (Johansson & Vallbo, 1979; Vallbo & Johansson, 1984). On the other hand, the hairy skin does not only possess myelinated mechanoreceptive afferents (with lower density; Provitera et al., 2007), but it also contains unmyelinated, slow conducting C-Tactile afferents (CTs; Vallbo et al., 1999; Vallbo, Olausson, Wessberg, & Norrsell, 1993). Specifically, microneurography studies on unmyelinated CT afferents suggest that CT afferents preferentially encode low force, relatively slow stroking touch. Namely, CTs activation follow an inverted U-shape relationship between the CT firing mean frequency rate and the stroking velocity; that is, CTs activation is higher in response to relatively slow velocity tactile stimulation (1-10cm/s^-1^) and lower in response to velocities above or below this range, suggesting that stroking within the 1-10 cm/s range optimally activates CT afferents. Moreover, such CT activation is strongly correlated with perceived pleasantness (Löken, Wessberg, Morrison, McGlone, & Olausson, 2009). No such relationship exists between the mean firing frequencies of myelinated afferents, such as for example Aβ fibres, and stroking velocity. Instead, the response of myelinated afferents increases with faster velocities, and shows no relationship with perceived pleasantness (Löken et al., 2009). Thus, the hypothesis has been put forwards that although the mammalian brain does have myelinated (Aβ) fibres that can encode tactile velocity faster than than unmyelinated fibres, it has a specialised CT afferent pathways in order to allow individuals to distinguish (‘pick up’) a range of velocities likely to have social-affective relevance for the purposes of further hedonic processing in affect-related brain areas such as the insula (Olausson et al., 2008). Thus, according to this so-called ‘social touch hypothesis’, gentle slow social touch is a distinct domain of touch, served by a specialised pathway capable of distinguishing affective information from the ‘‘noise’’ of tactile stimulation that does not carry any affective significance for the individual (see Morrison et al., 2010 for discussion).

Following the proposal of this hypothesis, a large body of research has focused on quantifying the sensory and emotional aspects of touch, including hairy and glabrous skin (Ackerley, Carlsson, Wester, Olausson, & Backlund Wasling, 2014; Ackerley, Saar, McGlone, & Backlund Wasling, 2014; Essick et al., 2010; Guest et al., 2011; Mcglone et al., 2012), and on conducting relevant neuroimaging studies to study the role of limbic and insular regions in the central processing of affective as opposed to neutral touch stimuli. For example, a growing body of research has characterised an affective touch network that appears to bypass the primary somatosensory area typically associated with the discriminatory aspects of touch S1 (c.f., Gazzola et al., 2012; Mccabe, Rolls, Bilderbeck, & McGlone, 2008) by activating brain regions implicated in the cognitive-affective aspects of the touch, such as the posterior superior temporal sulcus, medial prefrontal cortex orbitofrontal cortex (OFC) and anterior cingulate cortex (ACC) (Gordon et al., 2013). Specifically, although CT-optimal touch also activates the primary somatosensory cortex in healthy subjects, correlations with pleasantness ratings have only been reported in the insula and the OFC (Mccabe et al., 2008; Kress et al., 2011). Moreover, using fMRI and transcranial magnetic stimulation, evidence suggests that discrimination and intensity ratings predict activation in the S1, whereas pleasantness ratings predicted activation in the ACC (associated with the affective-motivational aspects of touch) but not the S1 (Case et al., 2016). Further, while investigating the cortical areas that represent affective touch, painful touch and neutral touch, studies have also found increased activity in the OFC in response to pleasant and painful touch, as compared to neutral touch, highlighting the role of the OFC on the affective aspects of the touch (Rolls et al., 2003). In contrast, the somatosensory cortex was less activated by pleasant and painful touch, relative to neutral touch (Rolls et al., 2003).

Important insights about the role of insular regions in the central processing of affective touch have been provided from patients with sensory neuronopathy (Olausson et al., 2002; Olausson et al., 2008). Given that sensory neuronopathy affects myelinated but not unmyelinated fibres, these patients are thought to lack Aβ afferents while their CTs afferents may remain intact. Interestingly, research has shown that CT stimulation in these patients activated the insula (i.e., the preferential cortical target for CT afferents; see more below), but not somatosensory regions associated with the sensory discriminative processing of touch (Olausson et al., 2002). On the other hand, these patients were able to detect, although poorly, slow brushing on the forearm (where CTs are abundant), which was accompanied by sympathetic skin response (Olausson et al., 2008). Given the sensory discriminative properties associated with Aβ-fibres and the lack thereof in these patients, it is possible to presume that while CT touch may not be involved in the discriminative aspects of touch, some somatotopical organization occurs in the insular cortex (Olausson et al., 2008; see also Bjornsdotter et al., 2009). Moreover, patients with hereditary sensory and autonomic neuropathy type V, i.e., presenting a reduction in density of thin and unmyelinated nerve, including CT afferents, perceived slow brushing on the forearm as less pleasant than controls, with such tactile stimulation not modulating activity in the posterior insula cortex – a target for CT afferents (Morrison, Bjornsdotter, & Olausson, 2011). Together, these findings suggest that CT afferents follow a separate neurophysiological route than Aβ mediated discriminative touch.

Despite this progress in neuroscientific studies, reliably assessing whether the CT system contributes to the ability to discriminate between affective and non-affective tactile stimuli has been hampered by the ongoing scientific challenges involved in quantifying the emotional perception of tactile stimuli. For example, Guest et al. (2011) developed and validated a descriptive scale for touch perception, namely the TPT, consisting of sensory and emotional descriptors. Using the TPT, it has been found that stroking the forearm (hairy, CT-skin) led to higher ratings of emotional descriptors, whereas stroking the palm (glabrous, non-CT skin) led to higher sensory descriptors (McGlone et al., 2012). Similarly, using different materials (i.e., soft brush, fur, sandpaper) has been shown to lead to higher emotional content on the forearm versus the palm and cheek (Ackerley, 2014). Nevertheless, the TPT has not been used to assess the emotional attributes of touch at different speeds, which as aforementioned, play an important role in both CTs firing rate and perceived pleasantness; perhaps because it would be difficult to use verbal descriptors when the only changing variable is the velocity of the touch. Instead, research on the field has mostly focused on the use of ratings, such as visual analogue scales on a continuous pleasantness dimension (e.g., ranging from not pleasant at all to extremely pleasant) and more recently evaluative conditioning (Pawling, Trotter, McGlone, & Walker, 2017).

Employing pleasantness ratings, numerous studies have shown that when stroking the forearm, individuals perceive gentle stroking touch within the CT-range, i.e., 1-10 cm/s, as more pleasant than gentle stroking touch outside the CT-range (e.g., Ackerley, Carlsson, et al., 2014; Ellingsen et al., 2014; I. Morrison, Bjornsdotter, & Olausson, 2011; Sehlstedt et al., 2016), showing a similar inverted U-shape relationship between stroking velocity and perceived pleasantness as reported by Löken et al (2009). However, the use of ratings in general has been subject to several criticisms, including individual differences in the use of the scale. Moreover, within the CT literature, the use of pleasantness ratings has not been able to provide a clear distinction between CT and non-CT skin. Indeed, given that no CTs have been found on glabrous skin, one would not expect an inverted U-shape relation between stroking velocities and perceived pleasantness on glabrous skin such as the palm. However, several studies report that stroking within a CT-optimal range also leads to similar increase in ratings of pleasantness in both forearm and palm (Ackerley, Carlsson, et al., 2014; Essick, James, & McGlone, 2009; Gentsch, Panagiotopoulou, & Fotopoulou, 2015; Löken, Evert, & Wessberg, 2011; Mcglone et al., 2012). Thus, it is thought that the effects of perceived pleasantness observed on the palm are likely due to secondary reinforcement, wherein top-down processes associated with expectations of pleasantness linked to relatively slow stroking can also lead to increased perception of pleasantness even when delivered to non-CT skin (McGlone et al., 2014). In this sense, pleasantness ratings may not be sufficient to discriminate between bottom-up sensory mechanisms (CT afferent signalling in response to touch at CT optimal speeds in CT skin) and top-down higher cognitive mechanisms (learned expectations linked to touch at CT optimal speeds) that may lead to the experience of pleasantness on the skin.

One way to tap into the sensory and cognitive domains that underpin the emotional aspects of touch is asking subjects to make forced decisions about the tactile stimuli, and assess the accuracy on their discrimination as well as their ‘attitudinal predisposition’. This way of assessing sensory and cognitive domains, based on ‘Signal Detection Theory’ (SDT), has been applied to both exteroceptive (e.g., vision, audition;, Cameron, Tai, Eckstein, & Carrasco, 2004; Hillyard, Squires, Bauer, & Lindsay, 1971; Verghese, 2001) and interoceptive modalities (e.g., heart beat activity, pain; Garfinkel et al., 2015; Mancini, Nash, Iannetti, & Haggard, 2014; Rollman, 1977) as a measure of exteroceptive or interoceptive sensitivity, respectively. Specifically, SDT provides a means by which one can measure an individual performance in detecting, or discriminating between, ambiguous stimuli by a known process (i.e., signal) or by chance (i.e., noise), providing two main outcomes: sensitivity (d’) and response bias (C), which mainly tap into the sensory and cognitive domain, respectively (Heeger, 2017; Macmillan, 2002; Rollman, 1977). Sensitivity (d’) indicates the strength of the signal (relative to the noise). Response bias (C) indicates the cognitive strategy or attitudinal tendency of the participant. These two parameters are extracted by calculating the proportion of trials in which the participant accurately detected the signal (i.e., hit rate) and the proportion of trials in which the participant said it was signal but it was in fact noise (false alarm rate) (see Abdi, 2007 for their respective formulas). Thus, the current study aimed to go beyond prior investigations of the social touch hypothesis by combining for the first time CT stimulation with signal detection theory (SDT) and examining whether the availability of CT afferents (in hairy versus in glabrous skin) will indeed lead to greater discrimination between tactile stimuli of high versus low affective relevance. In particular, consistent with the CT literature (Löken et al., 2009; McGlone et al., 2014), we predetermined the tactile stimuli administered at CT-optimal speeds (1-10 cm/s range) as more likely to generate ‘high pleasantness’ judgements (i.e., signal) and the stimuli administered at non-CT optimal speeds (slower than 1 cm/s and faster than 10 cm/s) as more likely to generate ‘low pleasantness’ responses (i.e., noise). We were thus able to assess the social ‘touch hypothesis’ by quantifying the ability of participants to accurately discriminate the touch in terms of pleasantness (high versus low), as well as their potential negative or positive bias (i.e., tendency to perceive the low-pleasant stimuli as high-pleasant or vice versa), in a forced-choice task on two different skin sites: forearm (CT-skin) and palm (non-CT skin) (type 1 task).

In addition, this study measured the participant’s own ability to recognize their trial-by-trial successful detection of pleasant stimuli (as high or low pleasant depending on CT versus non-CT optimal speeds, respectively) on forearm and palm. Specifically, this study employed a receiver operating characteristic (ROC) analysis (type 2 task) as a measure of metacognitive sensitivity (Fleming & Lau, 2014). This measure of metacognitive sensitivity has been applied to exteroceptive and interoceptive modalities (e.g., Fitzgerald, Arvaneh, & Dockree, 2017; Garfinkel et al., 2013, 2015; Palser, Fotopoulou, & Kilner, 2018) as an index of how participant’s confidence relates to their accuracy (type 2 hit) and inaccuracy (type 2 false alarm) on detecting a signal, while being ‘bias free’. For example, one may think that when one is confident, one is more likely to be correct. However, this is not always the case; one can have high overall confidence (e.g., a bias to provide high confidence ratings) that does not relate to correct responses, and vice-versa. Critically, although metacognitive measures have been applied to several modalities, with recent research arguing for the supramodality of metacognition, i.e., finding a correlation between visual, auditory, and tactile metacognitive efficiency (Faivre, Filevich, Solovey, Kühn, & Blanke, 2017), it still remains unknown whether these supramodal confidence findings extend to interoceptive signals, and whether any domain-specific processes also contribute to metacognition (Roualt et al., 2018). In this study, one could speculate differences in metacognitive sensitivity about affective touch on a skin site that possesses CT afferents versus one that does not. In particular, confidence can be regarded as a proxy for an individual’s subjective uncertainty over the accuracy of their perceptual decision. If CT-optimal touch is indeed a distinct, specialised modality for discriminating, affective touch as ‘the social touch hypothesis’ argues, then one would expect participants to be not only more accurate in discriminating between high and low tactile pleasantness in CT, hairy skin (level 1 performance) but also to be more confident about their discrimination accuracy in this location (level 2 performance), given the availability of more information about the affective relevance of the stimulus and a life-time of experience of such pleasantness discriminations. Thus, we anticipated that higher confidence would predicts pleasantness judgment’s accuracy better on the forearm than on the palm.

In sum, over two experiments, the present study aimed to test two major, experimental predictions stemming out of the wider ‘social touch hypothesis’ about the functional role of the CT system in affective touch discrimination. First, given that CTs are located on the forearm but not the palm, we hypothesized higher pleasantness sensitivity (d’) on the forearm versus palm, consistent with the specificity of bottom-up CT-based signalling. Second, we hypothesized higher confidence to predict the accuracy of detected pleasantness (as measured by the ROC analyses) better on the forearm versus the palm, given the availability of more information about the affective relevance of the stimulus on hairy skin and a life-time of experience of such pleasantness discriminations.

## 2. Experiment 1

### 2.1 Methods

#### 2.1.1 Participants

Based on an initial pilot using the exact same method outlined below, with a separate sample size of 21 participants, with an effect size of Cohen’s d= .27 and .30 for sensitivity d’ and response bias, respectively, power calculations indicated we would require at least ninety subjects to achieve a power of .80 (G*power 3.1). Thus, ninety-four participants (M_age_=22.97, SD_age_=5.56) were recruited via the University College London (UCL) Psychology Subject Pool for this study and received £10 or 1 credit in compensation for their time. Both females and males (Fourty-nine females, fourty-five males) were recruited. Even though the experimenter delivering the touch was always female, there were no gender effects across any of our outcome measures (see Supplementary Material). The UCL ethics committee approved this study and the experiment was conducted in accordance with the Declaration of Helsinki.

#### 2.1.2 Procedure

##### Signal Detection Preparation and Familiarisation Phase

Upon obtaining written informed consent, participants were informed that they would receive soft tactile stimuli on their arm and two adjacent stroking areas (each measuring 9 cm long) were marked on the participant’s forearm and palm (same arm). As an example of the kinds of touch they were about to receive in the main task, participants received one repetition of four randomised types of touch with a soft cosmetic brush (Natural hair Blush Brush, No. 7, The Boots Company; 1 stroke at 2 CT speeds: 3 cm/s and 9 cm/s; and 2 non-CT speeds: .5 cm/s and 18 cm/s). The velocity of the touch or their expected effects of felt pleasantness (i.e., high versus low) were not disclosed to participants. It’s worthwhile mentioning that we decided to not disclose the expected effects of pleasantness in the familiarisation phase because the expected effects could have driven the participants to base their judgements on this memory element and thus answer the main task based on these example velocities, irrespective of their actual perceived pleasantness.

##### Signal Detection Main Task

Participants were told they would receive similar kinds of touch on their forearm and palm, and they would have to focus on the sensation arising from the touch and categorise the touch into two categories of felt pleasantness: ‘high’ or ‘low’. In each trial, following the participant’s pleasantness judgement, they were also asked to provide confidence ratings on their judgment using a scale ranging from −4 ‘not at all confident’ to 4 ‘extremely confident’. Next, using the same soft cosmetic brush (Natural hair Blush Brush, No. 7, The Boots Company), the first half of the participants received 48 randomised touch trials of 4 CT-optimal speeds (1 cm/s, 3 cm/s, 6 cm/s, 9 cm/s) and 4 non-CT optimal speeds (0.3 cm/s, 0.5 cm/s, 18 cm/s, 30 cm/s; i.e., two velocities that were slower and two that were faster than the CT optimal range) on their forearm (CT-skin; 24 trials) and palm (non-CT skin; 24 trials). Interim analyses were conducted following this first half of the sample on our main outcome measures, i.e., sensitivity d’ and response bias (revealing the same pattern of results as those reported below with the total sample size; see supplementary materials). As there is a reasonably high chance to observe a significant effect after collecting only half of the participants suggested by a priori power analyses (Lakens, 2014), we continued testing for the planned total sample size but results below were interpreted under a lower alpha level set at .025 (i.e., alpha/2 given the two analyses: one interim and one after all the data was collected) to control for type 1 error as a result of performing interim analyses (see Lakens, 2014). Nevertheless, for the second half of the participants, we increased the number of touch trials to 64 randomised trials (i.e, 4 trials per velocity instead of 3), in order to increase reliability and approach a more normal distribution of what we predetermined as the signal and noise. This was accounted for in our calculation of d’ and C by adjusting the number of signal and noise trials (see below for d’ and C calculations). Moreover, independent sample t-tests conducted between the first and second half of the sample indicated no significant differences in any of our outcome measures, all *p*’s > .05, confirming that there were no differences on d’, C and aROC between the first and second half of our sample.

##### Manipulation check (pleasantness ratings)

Following the main Signal Detection task, we collected pleasantness ratings of CT optimal (1 cm/s, 3 cm/s, 6 cm/s, 9 cm/s) and non-CT optimal (0.3 cm/s, 0.5 cm/s, 18 cm/s, 30 cm/s) speeds in order to make sure that participants were in fact perceiving touch at CT-optimal speeds as more pleasant than non-CT optimal speeds (this is important for SDT analyses given we are considering CT-optimal speeds as the signal). One trial per velocity was delivered at each skin site (with the order randomised across skin site). We used the same soft brush to administer the randomized 16 trials. After each trial, participants were instructed to rate verbally the pleasantness of the touch using a scale ranging from 0 ‘not at all pleasant’ to 100 ‘extremely pleasant.

##### Tactile acuity

Following the control task, general tactile sensitivity to punctuate touch was also tested using Von Frey monofilaments in a force detection paradigm (only on the second half of the sample, i.e., n=47, consistent with prior studies employing a similar sample size finding significant results when comparing tactile acuity on forearm versus palm; e.g., Ackerley et al., 2014). As determined in pre-tests, five calibrated monofilaments with sufficient range of forces were chosen: 0.4 mN, 0.7 mN, 1.6 mN, 3.9 mN, 9.8 mN, 19.6 mN, 39.2 mN. Using an increasing/decreasing detection difficulty task (Bell-Krotoski et al., 1993), we established a threshold monofilament for each skin site (i.e., forearm and palm; the order of skin site was counterbalanced across participants). Each monofilament was pressed five times (for 1s with a 1s gap) and after the five presses, the participant was asked to say how many presses they felt.

#### 2.1.3 Data analyses

##### Signal detection (sensitivity d’ and response bias C)

In order to implement signal detection analysis, and consistent with prior literature on CT stimulation and perceived pleasantness (Löken, et al., 2009), the stimuli administered at CT-optimal speeds (1 cm/s, 3 cm/s, 6 cm/s, 9 cm/s) were considered as more likely to generate ‘high pleasantness’ judgements, i.e., signal, whereas the stimuli administered at non-CT optimal speeds were considered as more likely to generate ‘low pleasantness’ responses, i.e., noise (see Figure 1). We thus calculated normalised hit rates (P[‘‘high’’/CT-optimal speed], i.e. the proportion of hit trials to which subject responded ‘‘high’’ and the stimulus was administered at CT-optimal speeds), and false alarm rates (P[‘‘high’’/non-CT optimal speeds], i.e. the proportion of trials in which the touch was administered at non-CT optimal speeds but the participant responded ‘‘high’’) for the forearm (CT skin) and palm (non-CT skin) separately. These were used to obtain the perceptual sensitivity (d’), a measure of discriminability in detecting the high-pleasure target, and the response bias (C), which measures the tendency to report stimuli as ‘‘high’’. The *affective touch sensitivity* (d’) was quantified as: d’ = z(hit rate)-z(false alarm rate). The response bias (C) can be expressed as: C = (z[hit rate] + z[false alarm rate]) * 0.05. Data from one participant was not included given that the number of ‘high’ responses to low-pleasant stimuli was equivalent to the maximum (i.e., 16 ‘high pleasant’ responses were given to the 16 stimuli predetermined to generate ‘low pleasantness’ responses, i.e., noise), and thus d’ and C could not be computed. This yielded a total sample size of 93 participants for these analyses. We conducted paired t-tests between forearm and palm separately on sensitivity (d’) and response bias (C), to examine whether there was any difference in sensitivity (d’) and response bias (C) between CT and non-CT skin sites.

**Figure 1:**
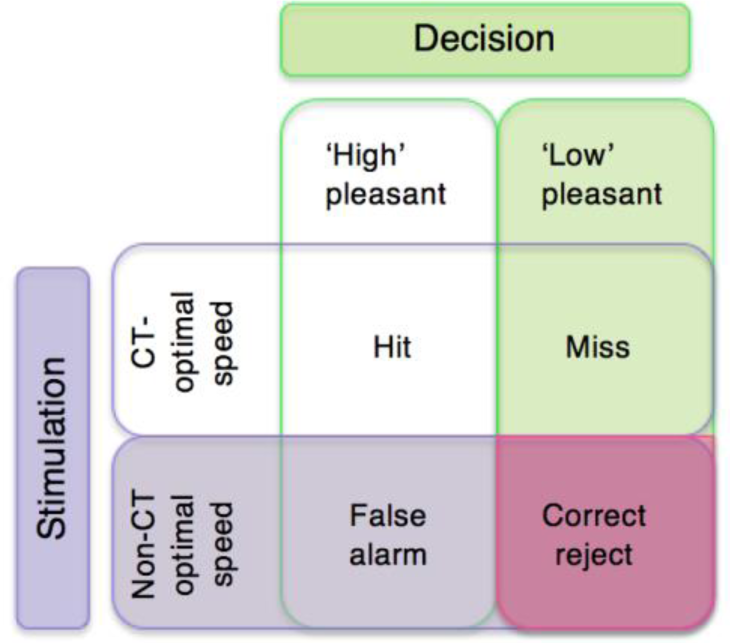
A schematic representation of the SDT model applied to affective touch. CT-optimal speeds include 1 cm/s, 3 cm/s, 6 cm/s, 9 cm/s, which were considered as the high-pleasant stimuli. Non-CT optimal speeds include 0.3 cm/s, 0.5 cm/s, 18 cm/s, 30 cm/s; i.e., two velocities that were slower and two that were faster than the CT optimal range) and were considered as the low-pleasant stimuli. Normalised hit rates were calculated as (P[‘‘high’’/CT-optimal speed], i.e. the proportion of hit trials to which subject responded ‘‘high’’ and the stimulus was administered at CT-optimal speeds), and false alarm rates were calculated as (P[‘‘high’’/non-CT optimal speeds], i.e. the proportion of trials in which the touch was administered at non-CT optimal speeds but the participant responded ‘‘high’’).

##### Metacognitive sensitivity

To investigate the extent to which confidence predicts perceptual accuracy, i.e., correct and incorrect responses about the categorization of the tactile stimuli in terms of pleasantness and speed (i.e., type 1, high/low-pleasant, see above), a type 2 receiver operating characteristic (ROC) analysis was conducted. ROC examines the extent to which a signal (here confidence) is an effective detector of some binary variable *i*. For each confidence rating threshold (in this case 10), one computes the hit rate (i.e., proportion of trials correctly detected, e.g., proportion of *i*=1 trials detected as *i*=1; simply put, proportion of correct trials in which confidence was ‘high’) and the false alarm (proportion of trials incorrectly detected; e.g., *i*=0 trials detected as *i*=1; simply put, proportion of incorrect trials for which the confidence was ‘high’). The ROC curve plots the hit rate versus the false alarm rate over all the possible detection thresholds. To measure how confidence is predictive of accuracy one typically computes the area under the ROC curve (aROC). The aROC provides a measure of the extent to which the detector is correct for a given level of confidence (Massoni, Gajdos, & Vergnaud, 2014) and therefore can be used as a measure of metacognitive sensitivity. Higher aROC indicates higher metacognitive sensitivity (Fleming & Lau, 2014). Data from thirty-six participants was not included in this analysis, given that there was not enough variance in the confidence ratings between the binary variable *i* for these participants and thus an aROC could not be computed (yielding a sample of 58 participants for this analysis). We conducted paired t-tests between forearm and palm separately on aROC scores, to examine whether higher confidence predicts accuracy better on CT versus non-CT skin.

##### Manipulation check (pleasantness ratings)

CT optimal (1 cm/s, 3 cm/s, 6 cm/s, 9 cm/s) and non-CT optimal (0.3 cm/s, 0.5 cm/s, 18 cm/s, 30 cm/s) touch ratings were averaged separately per skin area (forearm, palm) for each participant, creating CT-optimal touch palm, CT-optimal touch forearm, non-CT optimal palm and non-CT optimal forearm pleasantness averaged rating scores for each participant (see supplementary materials for group level analyses on pleasantness scores). To further validate our signal detection model in the main task, correlations were conducted between the sensitivity (d’) scores of our participants and their difference scores on the averaged subjective pleasantness ratings (CT-optimal speed minus CT non-optimal speed) separately on forearm and palm.

##### Tactile Acuity

The threshold level for each participant (per skin site) was defined as the monofilament at which the participant could feel at least four (out of five) presses in both the increasing and decreasing detection difficulty aspects of the task. Von Frey filaments were then dummy coded from 1 to 7, from lower to higher filament force (i.e., 0.4 mN=1, 0.7 mN=2, 1.6 mN=3, 3.9 mN=4, 9.8 mN=5, 19.6 mN=6, 39.2 mN=7) for the purpose of this analyses. Data from 2 participants were removed as they reported more than five presses on either the ascending or descending order, yielding a total sample size of 45 participants for this analysis. In order to make sure participant’s ability to discriminate pleasantness better between skin sites did not depend merely on general tactile sensitivity, we conducted correlations, *r*, between d’ and the threshold level on both skin sites. We also conducted paired t-tests on the threshold level between forearm and palm to assess whether there was more tactile sensitivity on palm versus forearm. On a scale from 1-7, lower scores would indicate greater tactile sensitivity.

#### 2.2 Results

##### 2.2.1 Affective Touch Sensitivity (d’) and Response bias (C)

As expected, significantly higher sensitivity (d’) was found on the forearm (*M*=1.43, *SD*=.91), as compared to the palm (*M*=1.19, *SD*=.94), *t*(92)=2.80, *p*=.006, *d*=0.29 (see Figure 2A), even at an alpha level set at .025 (see section 2.1.2). However, surprisingly we also found more negative response bias (C) on the forearm (M= -.32, SD=.47) as compared to the palm, (M= -.14, SD=.56), t(92)=2.76, *p*=.007, *d*=0.29 (see Figure 2B; bottom panel), even at an alpha level set at.025 (see section 2.1.2). Thus, these results suggest that participants are better at discriminating high-pleasure target stimuli on the forearm vs. palm, with a tendency to also report low-pleasant stimuli as high pleasant on this same skin site.

**Figure 2.**
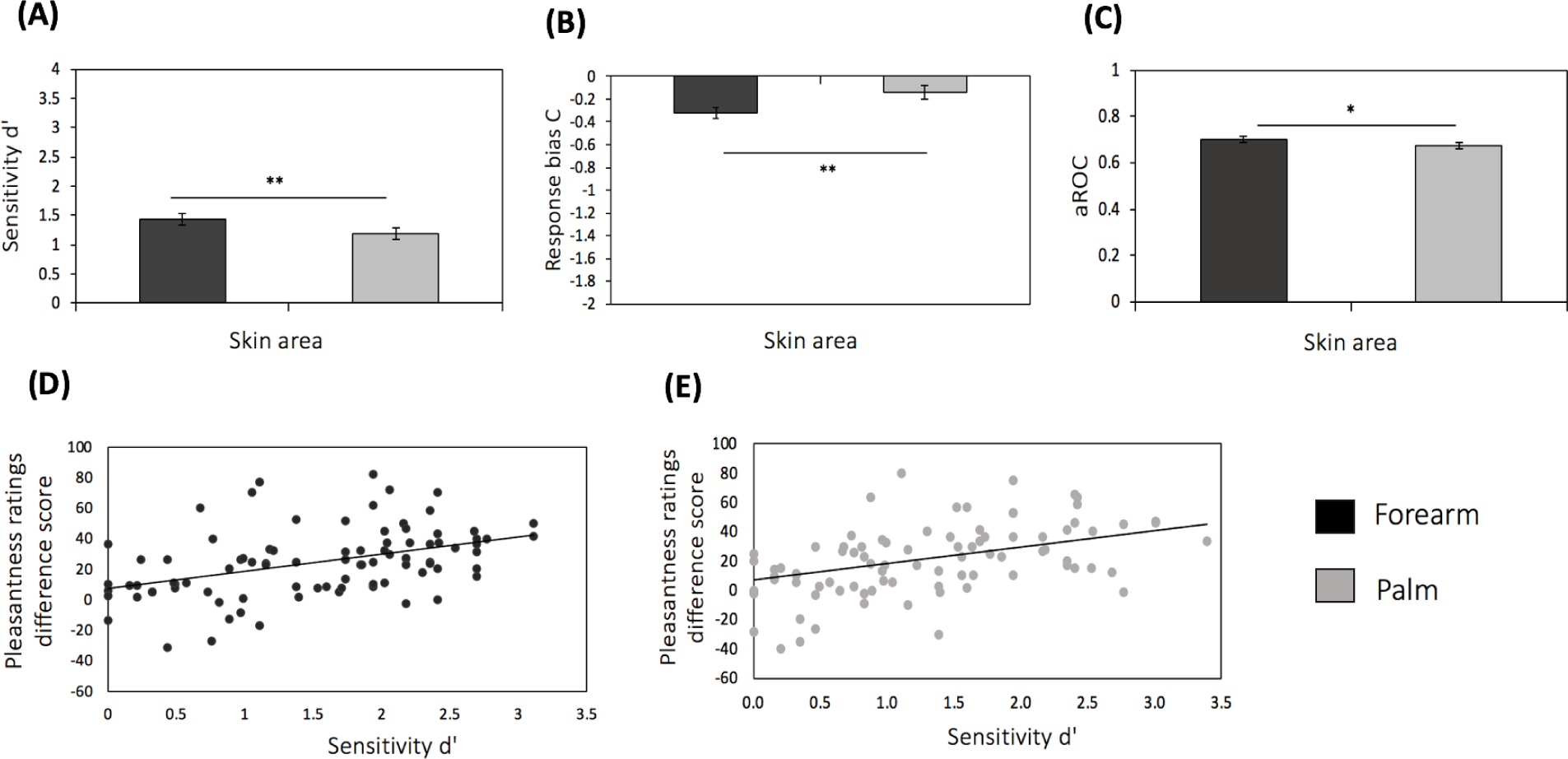
Experiment 1 Results. **(A)** Sensitivity (d’) for forearm and palm. **(B)** Response bias C for forearm and palm. **(C)** aROC for forearm and palm. **(D)** Correlation between sensitivity d’ (x-axis) and pleasantness ratings difference score (CT-optimal – non-CT optimal speed; y-axis) on the forearm. **(E)** Correlation between sensitivity d’ (x-axis) and pleasantness ratings difference score (CT-optimal – non-CT optimal speed; y-axis) on the palm. Error bars denote ± standard error of the mean for illustration purposes. * indicate statistical significance, p<.05. ** indicate statistical significance, p<.01

##### 2.2.2 Does confidence predict accuracy?

Our aROC analysis measured the extent to which confidence was predictive of the success in correctly categorizing the touch as high or low pleasant (hit) than in inaccurate responses (false alarm), as a measure of metacognitive sensitivity, separately on the forearm and the palm. We found greater metacognitive sensitivity on the forearm as compared to the palm, i.e., greater area under the ROC curve was found on the forearm (M=.71, SD=.12), relative to the palm (M=.67, SD=.11), t(57) =2.12, *p* = .038, *d*=0.28 (see Figure 2C).

##### 2.2.3 Manipulation check (pleasantness ratings)

Analyses conducted on the pleasantness ratings scores suggested that CT-optimal touch (at CT speeds; 1 cm/s, 3 cm/s, 6 cm/s, 9 cm/s) was perceived as more pleasant than non-CT optimal touch (at non-CT optimal speeds; 0.3 cm/s, 0.5 cm/s, 18 cm/s, 30 cm/s) at the group level (see Supplementary Materials). Further validating our signal detection model in the main task, we report a positive correlation between sensitivity d’ and the pleasantness rating difference scores (CT-optimal speed – non-CT optimal speed) on the forearm, r=.45, p<.001, and palm, r=.46, p<.001 (see Figure 2D and 2E). Thus, the bigger the difference on pleasantness ratings between CT and non-CT speeds, the higher their d’ scores, on both forearm and palm.

##### 2.2.4 Tactile acuity

As expected, tactile acuity sensitivity as measured by the von Frey filaments differed across skin sites, t(44) =11.89, *p*<.001, *d*=1.77. Specifically, higher tactile sensitivity was observed on the palm (M=2.33, SD=1.00) versus the forearm (M=4.44, SD=1.03). Moreover, tactile sensitivity did not correlate with d’ sensitivity on the palm, r= -.06, p=.694, nor forearm, p= -.15, p=.330, indicating that participant’s ability to discriminate pleasantness better between skin sites did not depend merely on general tactile, acuity sensitivity.

##### 2.2.5 Summary of results for Experiment 1

In sum, consistent with our predictions and the social touch hypothesis, these results showed higher pleasantness (d’) sensitivity on the forearm (CT skin) versus the palm (non-CT skin), even though higher tactile acuity as measured by the von Frey filaments was observed on the palm versus the forearm. In other terms, the touch delivered at CT-optimal speeds was judged as ‘high-pleasant stimuli’ (i.e., hits) more frequently than touch delivered at non-CT optimal speeds (i.e., less false alarms) in a skin area that contains CT afferents, i.e. the forearm, in comparison with a skin area that does not contain CT fibres, i.e the palm. By contrast, as a measure of tactile acuity, participants were able to detect to a better extent the von Frey stimuli (i.e., static, brief tactile stimuli) on a skin site that does not possess CT afferents, i.e. the palm, relative to the forearm. Similarly, higher confidence was more predictive of accurately detecting pleasantness on the forearm versus the palm, suggesting better metacognitive sensitivity on CT-skin relative to non-CT skin (i.e., higher confidence in accurate [hit] relative to inaccurate responses [false alarm]). However, surprisingly, we also observed more negative response bias (i.e., more hits and false alarms) on the forearm versus the palm, indicating a bigger tendency or ‘attitudinal predisposition’ to generally judge tactile stimuli as high-pleasant when administered on CT skin, irrespective of their speed. Finally, as a manipulation check, we collected pleasantness ratings from each stroking velocity on both forearm and palm. Positive correlations between pleasantness ratings difference score (the average of CT speeds minus the average non-CT speeds, termed hereafter ‘CT ratings differential’) and pleasantness (d’) sensitivity indicated a moderate-to-strong positive relationship between d’ and CT ratings differential on both forearm and palm. This relationship further validates our signal detection model in the main task by indicating that pleasantness (d’) sensitivity, which is based on a binary, discriminatory response of high versus low pleasantness captures pleasantness perception in a similar way as widely-used, pleasantness ratings scales.

Nevertheless, there is the possibility that some of these results can be explained by order effects of skin site associated with the perception of pleasantness following tactile stimulation on a skin site that contains CT-afferents. In particular, recent evidence suggests that the perception of pleasantness for palm stimulation is influenced by preceding forearm stimulation. It has been suggested that there is not only a general increase in perceived pleasantness following tactile stimulation on the forearm (which contains CT afferents) but also an increased pleasantness in response to touch at CT-optimal versus non CT-optimal speeds on this non-CT skin area (Löken et al., 2011). Thus, one may speculate that in a skin area where there are more top-down biases about CT-optimal velocities, such as the palm, the felt pleasantness following touch in the forearm may ‘spill over’ to this non-CT innervated skin area (and even to speeds outside the CT range). Indeed, results from Experiment 1 further suggest there is also a negative response bias in the palm (and not just the forearm), reflecting a tendency to judge the touch as ‘high-pleasant’. Accordingly, a block design was employed to measure signal detection separately on each skin site in Experiment 2.

## 3. Experiment 2

### 3.1 Methods

#### 3.1.1 Participants

Based on the initial pilot (see section 2.1.1) and given Experiment 1 (also revealing similar effect sizes), suggesting that at least ninety subjects are needed to obtain a power of .80, a similar sample size was selected for this experiment. Specifically, one hundred participants (M_age_=21.9, SD_age_=3.34) were recruited via the University College London (UCL) Psychology Subject Pool. The study was conducted in accordance with the Declaration of Helsinki and was approved by the University’s ethics committee and participants received £10 or 1 credit in compensation for their time. Distinct to Experiment 1, in this experiment a block design was employed to measure signal detection separately on each skin site (i.e., the touch trials from the main SDT task were delivered first on the palm and then on the forearm, with the order counterbalanced across participants). Specifically, forty-nine participants were assigned to group A, in which they first completed the Signal Detection task on their forearm (block 1), and then completed the Signal Detection task on their palm (block 2). Conversely, fifty-one participants were assigned to group B, in which they first completed the Signal Detection only on their palm (block 1), and then completed the Signal Detection task on their forearm (block 2). Females and males were recruited (fifty-three females: twenty-five assigned to group A and twenty-eight to group B; forty-seven males: twenty-four assigned to group A and twenty-three to group B). Similar to Experiment 1, even though the experimenter delivering the touch was female, there were no gender effects across any of our outcome measures (see Supplementary Materials).

#### 3.1.2 Procedure

Upon obtaining written informed consent, participants were informed that they would receive soft tactile stimuli on their arm and two adjacent stroking areas (each measuring 9 cm long) were marked on the participant’s forearm and palm (same arm). As mentioned above, participants completed both the Signal Detection Task separately on their forearm and palm, with the order counterbalanced across participants (i.e., group A or B). For example, on the Signal Detection task – forearm block, participants were given the same instructions as described in Experiment 1 (see section2.1.2 ‘signal detection preparation and familiarization phase’). This was followed by the Main Signal Detection task – Forearm, where participants received 32 touch trials (4 trials per velocity, same velocities described in Experiment 1) but only on their forearm and asked to categorise the touch into two categories of felt pleasantness: ‘high’ or ‘low’, followed by confidence ratings on their judgment using a scale ranging from −4 ‘not at all confident’ to 4 ‘extremely confident’ (as in Experiment 1).

Following the completion of these two blocks, participants completed the same manipulation check described in Experiment 1 (i.e., pleasantness ratings were collected for each of the eight velocities; 4 CT and 4 non-CT). The order of skin site in this last control task resembled the order of blocks to which each participant had been assigned.

#### 3.1.3 Data analyses

##### Signal detection (sensitivity d’ and response bias C)

Sensitivity d’ and C for the forearm and palm was calculated in the same way as in Experiment 1 (see section 2.1.3 for details). Here, data from six participants were not included given that the number of ‘high’ responses to low-pleasant stimuli was equivalent to the maximum on either their forearm or palm (i.e., 16 ‘high pleasant’ responses were given to the 16 stimuli predetermined to generate ‘low pleasantness’ responses, i.e., noise), and thus d’ and C could not be computed. This yielded a total sample size of 94 participants for these analyses. We conducted a 2×2 mixed ANOVA, specifying order (first forearm, first palm) as between-group factor, separately on sensitivity (d’) and response bias (C), to examine whether there was any difference in sensitivity (d’) and response bias (C) between CT and non-CT skin sites, while examining potential order effects. Follow-up analysis used paired or independent t-tests with Bonferroni correction when applicable.

##### Metacognitive sensitivity

The aROC for the forearm and palm was calculated in the same way as in Experiment 1 (see section 2.1.3). Data from twenty-six participants was not included in this analysis, given that there was not enough variance in the confidence ratings between the binary variable *i* for these participants and thus an aROC could not be computed (yielding a sample of 74 participants for this analysis). We conducted a 2 (skin site: forearm, palm) x 2 (order: firs palm, first forearm) mixed ANOVA on aROC scores to examine whether high confidence predicts accuracy (hits) versus inaccuracy (false alarms) better on CT versus non-CT skin and whether such effects depend on the order of the block.

##### Manipulation check (pleasantness ratings)

We analysed pleasantness ratings from our control task as described in Experiment 1 (see section 2.1.3 for analyses details).

### 3.2 Results

#### 3.2.1 Affective Touch Sensitivity (d’) and Response bias (C)

Similar to Experiment 1, we found higher d’ on the forearm (M=1.74, SD=.99) than on the palm (M=1.38, SD=1.01), *F*(1,93)=10.93, *p*=.001, η^2^_partial_=.11. There was no main effect of order *F*(1,93)=.39, *p*=.573, η^2^_partial_=.00, and order did not interact with skin area, *F*(1,93)=.16, *p*=.693, η^2^_partial_=.00 (see Figure 3A). These results suggest that participants are better at discriminating high-pleasure target stimuli on the forearm vs. palm, irrespective of the order of the blocks.

**Figure 3.**
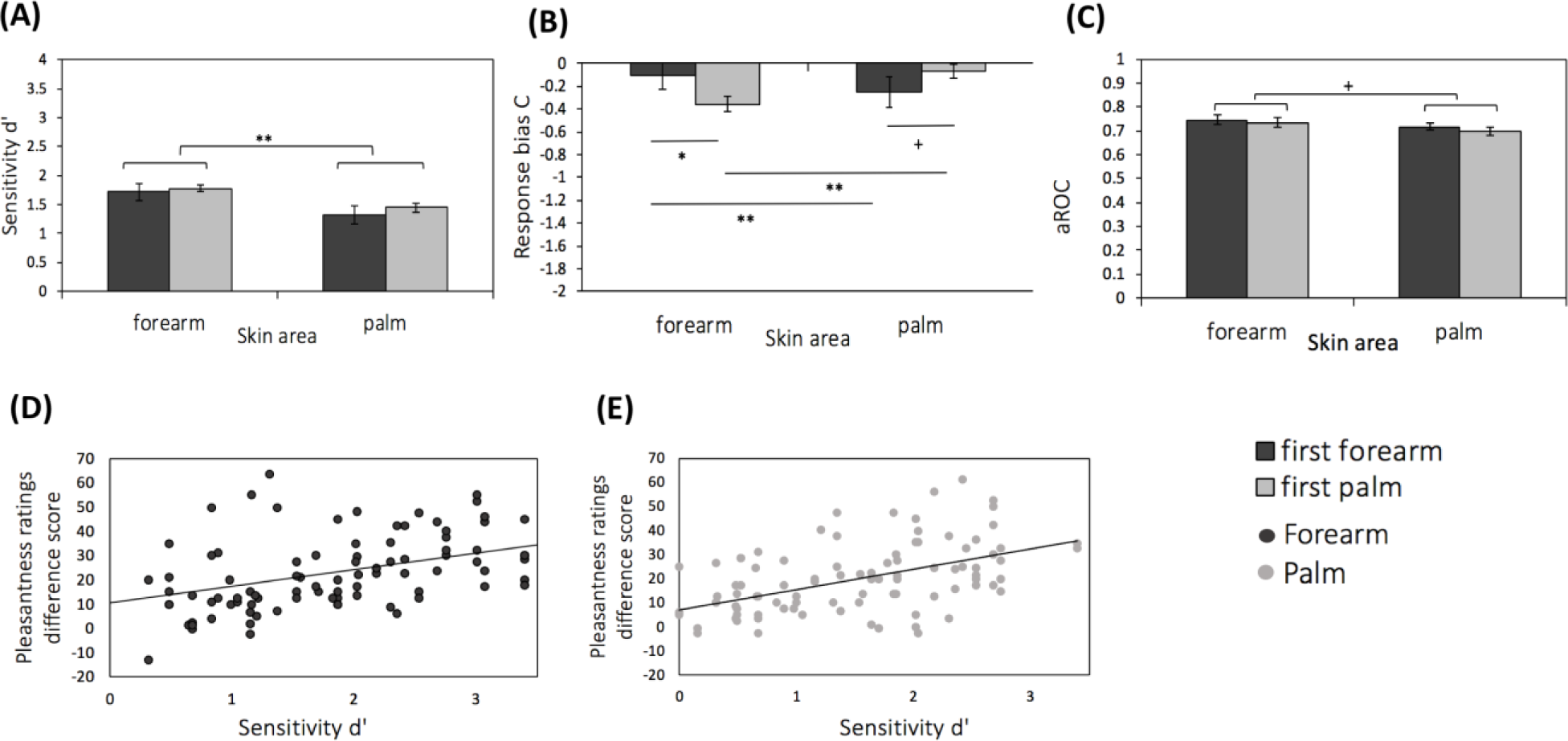
Experiment 1 Results. **(A)** Sensitivity (d’) for forearm and palm. **(B)** Response bias C for forearm and palm by order of the blocks. **(C)** aROC for forearm and palm by order of the blocks. **(D)** Correlation between sensitivity d’ (x-axis) and pleasantness ratings difference score (CT-optimal – non-CT optimal speed; y-axis) on the forearm. **(E)** Correlation between sensitivity d’ (x-axis) and pleasantness ratings difference score (CT-optimal – non-CT optimal speed; y-axis) on the palm. Error bars denote ± standard error of the mean for illustration purposes. * indicate statistical significance, p<.05. ** indicate statistical significance, p<.01

With respect to response bias (C), there was no main effect of skin area, *F*(1,93)=2.16, *p*=.145, η^2^_partial_=.02, or order, *F*(1,93)=.08, *p*=.794, η^2^_partial_=.00. However, skin area interacted with order, *F*(1,93)=27.28, *p*=<.001, η^2^_partial_=.23. Post-hoc tests, using Bonferroni adjusted alpha level of.0125 per test (.05/4), showed that if participants completed the palm block first, then there was more negative response bias on the second block, i.e., forearm, *t*(47)=4.28, *p*<.001, *d*=0.62. Similarly, if they completed the forearm block first, then there was more negative response bias on the second block, i.e., palm, *t*(46)=3.04, *p*=.004, *d*=0.44. However, between-subjects analyses showed that when the palm versus the forearm goes first, there was more negative response bias on the forearm, *t*(93)=2.60, *p*=.011, *d*=0.27, but not the palm, *t*(93)=1.94, *p*=.055, *d*=0.20 (see Figure 3B). These results suggest that exposure to touch may be associated with a general tendency to judge the touch as ‘high pleasant’, irrespective of skin area (although between-subjects effects suggest that such effects are more prominent if the palm is stimulated first), as reflected by more negative bias on the second block.

#### 3.2.2 Does confidence predict accuracy?

Similar to Experiment 1, although only at trend level, we found that high confidence was more predictive of accurately detecting pleasantness on the forearm versus the palm, i.e., greater area under the ROC curve was found on the forearm (M=.74, SD=.14) relative to the palm (M=.71, SD=.11), *F*(1,73)=2.34, *p*=.131, η^2^_partial_=.03, thus hinting to greater metacognitive sensitivity on CT-skin. There was no main effect of order, *F*(1,73)=.69, *p*=.408, η^2^_partial_=.00, and order did not interact with skin area, *F*(1,73)=.17, *p*=.684, η^2^_partial_=.00 (see Figure 3C).

#### 3.2.3 Manipulation check (pleasantness ratings)

Similar to Experiment 1, analyses conducted on the pleasantness ratings scores suggested that CT-optimal touch (at CT speeds; 1 cm/s, 3 cm/s, 6 cm/s, 9 cm/s) was perceived as more pleasant than non-CT optimal touch (at non-CT optimal speeds; 0.3 cm/s, 0.5 cm/s, 18 cm/s, 30 cm/s) at the group level (see Supplementary Material). Further validating our signal detection model in the main task, we report a positive correlation between sensitivity d’ and the pleasantness rating difference scores (CT-optimal speed – non-CT optimal speed) on the forearm, r=.44, p<.001, and palm, r=.59, p<.001 (see Figure 3D and 3E). Thus, the bigger the difference on pleasantness ratings between CT and non-CT speeds, the higher the d’ scores, on both forearm and palm.

#### 3.2.4 Summary of results for Experiment 2

In sum, consistent with Experiment 1, results from Experiment 2 showed higher d’ sensitivity on the forearm (CT skin) versus the palm (non-CT skin), irrespective of order effects. Our effects on criterion C, suggest that the tendency to judge the touch as high-pleasant stimuli depends on the order of the block, with more exposure to touch being associated with a bigger tendency to report the stimuli as high pleasant, irrespective of skin area. Our aROC analyses show a similar pattern for metacognitive sensitivity as reported in Experiment 1, although such effects are only present at trend level in Experiment 2. Finally, analyses conducted on the pleasantness ratings further validated our signal detection model in the main task by indicating a moderate-to-strong positive relationship between d’ and pleasantness ratings differential on both forearm and palm (as also seen in Experiment 1).

## 4. Discussion

The social touch hypothesis posits that gentle slow touch is a distinct domain of touch, served by the specialized CT pathway capable of distinguishing affective information from the “noise” of tactile stimulation that does not carry affective significance for the individual. However, previous research on this hypothesis has mostly relied on asking subjects to rate the perceived pleasantness of tactile stimuli using measures such as visual analogue scales or Likert-type scales that are of limited validity when it comes to determining discrimination abilities and more generally are subject to individual response tendencies and biases, such as individual differences in the interpretation and use of the scale. Here we used a forced-choice task that asks for a binary response, i.e., to judge the touch as high- or low-pleasant, with an analysis approach that can take into account individual response tendencies. Specifically, by combining for the first time CT stimulation with a signal detection task (SDT), we extracted different parameters, such as sensitivity d’, response bias, and metacognitive sensitivity (aROC), to provide an index of affective touch sensitivity and awareness. In Experiment 1, we implemented SDT with the touch trials randomized across CT and non-CT skin sites, i.e., forearm and palm, respectively. In Experiment 2, we implemented SDT using a block design (first forearm and then palm and vice versa). Supporting the social touch hypothesis and our first prediction, in both experiments, we found higher sensitivity d’ on the forearm versus the palm, indicating that people are better at discriminating high-pleasure target stimuli (i.e., touch at CT-optimal speeds) on a skin site that contains CT afferents. Surprisingly however, we found more negative response bias (C) on the forearm versus the palm, indicating a tendency to report the stimuli as ‘high-pleasant’ on CT-skin, irrespective of the touch velocity (i.e., CT-optimal and non-CT-optimal speeds) (Experiment 1). Nevertheless, in a follow-up experiment we assessed a potential explanation for this finding, namely that such effects could be driven by order effects, and indeed we found that there is more negative bias following touch exposure, but irrespective of skin site (Experiment 2). Supporting our second prediction, we found that there was more metacognitive sensitivity on the forearm than the palm, i.e., high confidence was more predictive of perceived (pleasantness) accuracy on the forearm versus the palm (although such effects were found only at trend level in Experiment 2). Further validating our SDT model, we found a moderate-to-strong relationship between d’ and pleasantness ratings differential (obtained from the manipulation check) on both forearm and palm in both experiments. These findings are discussed in turn below.

Microneurography studies have shown that the firing frequency of CTs, which have been found on hairy but not glabrous skin, and the stroking velocity follow an inverted U-shape relationship – with firing frequency of CTs being also highly correlated with pleasantness ratings (Loken et al., 2009). Such finding thereby suggests that there is a relationship between perceived pleasantness and coding at a peripheral level, hence perceived pleasantness on hairy skin is, at least partly, mediated by bottom-up CT afferent signalling. Beyond a correlational approach, however, the social touch hypothesis posits that CT afferents work as selectors to “pick out” a range of velocities likely to have social-affective relevance (Morrison et al., 2010). The present findings on sensitivity d’ lend support to this hypothesis by showing better discrimination to CT-optimal speeds (i.e., stimuli predetermined as ‘signal’), on the forearm versus the palm, even though there was better tactile acuity on the palm versus the forearm as measured by the von Frey filaments. The latter finding was expected given that glabrous skin in the hand is known to contain dense myelinated tactile afferents (Vallbo & Johansson, 1984) that send fast, temporally-accurate information about the touch to the brain, explaining the high tactile acuity. Similar findings on tactile acuity on glabrous versus hairy skin, including skin areas such as the palm and the forearm, have been shown by Ackerley et al. (2014). In fact, it has been proposed that the distribution of afferents on these skin sites reflects the usage of these skin surfaces, with for example the palm of the hands being key in exploring our environment (where a larger distribution of Aβ myelinated afferents exists) and skin sites such as the forearm or face being key for interpersonal interactions (where a lower distribution of Aβ myelinated afferents exists yet these skin sites also contain unmyelinated CT afferents) (see Ackerley, 2014; also see Vallbo & Johansson, 1984; Johansson et al., 1988; Vallbo et al., 1995).

Returning to our findings on sensitivity d’, it is worth noticing that the sensitivity d’ found in both forearm and palm is above chance. In fact, when it comes to sensitivity d’, zero values indicate no discrimination, but values above this range indicate at least some discrimination between these the stimuli predetermined as noise and signal – with the larger the d’ value, the better the discrimination. In practice, d’ of 4 or more indicates nearly perfect performance, and particularly in our study, given the number of signal and noise trials, d’ of 3.725 indicates nearly perfect performance. Thus, participants in the present study were also accurately determining, to a certain degree, CT-optimal speeds on the palm (i.e., stimuli predetermined as ‘signal’) as ‘high-pleasant’ stimuli, and stroking velocities outside the CT range (i.e., stimuli predetermined as ‘noise’) as ‘low-pleasant’ stimuli (where no CTs have been found). On the one hand, these findings also raise the possibility that even if the CT system is somewhat specialised for affective touch, it is not the only afferent system contributing to tactile pleasantness and further work is needed to establish how the various tactile afferent systems interact to give rise to feelings of tactile pleasantness in different body parts. For example, considering the afferent and efferent connections between posterior insular regions and parietal somatosensory regions (Augustine, 1996), it is likely that affective coding of tactile stimuli in the insular context may modulate somatosensory responses in SI (Olausson et al., 2008) and vice versa (see also Morrison et al., 2010). On the other hand, it is possible that such discrimination on the palm is a result of carry-over effects from preceding tactile stimulation from the forearm (CT skin). Indeed, previous research has shown that the perception of pleasantness on the palm is affected by stimulation on the forearm (Löken et al., 2011). However, we think this is unlikely because, as shown in Experiment 2, the order of the blocks (e.g., completing the SDT task first on the forearm and then on the palm) had no effects on sensitivity d’. Instead, our findings on sensitivity d’ suggest that this non-CT skin sensitivity is not affected by short, previous stimulation on CT-skin. We therefore speculate that such effects on the palm are due to learned associations between stroking velocity and pleasantness, which have been well reinforced throughout the lifespan. The precise role of CT system in such learning process still needs to be examined.

Turning to our second main finding, we unexpectedly found that there is more negative response bias (C) on the forearm versus the palm (resulting in more ‘hits’ and ‘false alarms’ in the former), indicating a bias to report the stimuli as ‘high-pleasant’ on CT-skin, irrespective of the touch velocity (Experiment 1). Such finding is consistent with previous research suggesting that CT skin, such as the forearm, is typically perceived as more pleasant than non-CT skin, such as the palm (e.g., Löken et al., 2011; Ackerley et al., 2014, except for speeds at 30 cm/s; see also supplementary materials for pleasantness ratings manipulation checks). Critically, however, a negative response bias as reflected by negative values (C<0) was found on both the forearm and palm, indicating a tendency to report the stimuli as ‘high-pleasant’ irrespective of touch velocity in both CT and non-CT skin sites. These results may suggest that there are top-down effects of perceived pleasantness on both forearm and palm – though to a larger extent on the forearm. In line with the social touch hypothesis (Morrison et al., 2010), we speculate that the tendency to judge the touch as high versus low pleasant is associated with the exposure to touch, which may be greater in skin areas that are key for social interactions such as the forearm. In fact, recent evidence suggests that people with less touch exposure show a general tendency to rate gentle, stroking touch as less pleasant (Sailer & Ackerley, 2017). Supportive of this notion, our findings in Experiment 2 further suggest that the observed tendency to judge the tactile stimuli as high-pleasant stimuli depends on the order of the block, with more exposure to touch being associated with a larger tendency to report the stimuli as high pleasant, irrespective of skin area. Future work is needed to investigate if this is the case. As alternative explanations, it is also possible that participants may be reluctant to categorise the touch as ‘low-pleasant’, which indeed has been a focus of recent debate regarding binary, similar ‘yes-no’ tasks as participants show substantial criterion effects (see Ruby, Giles, & Lau, preprint for a discussion) and thus, two-alternative-forced choice paradigm might be more optimal, particularly given that gentle, strokes by an experimenter are particularly unlikely to be judged as unpleasant and indeed can be thought of as carrying some socio-affective value. Regarding our second hypothesis, our aROC findings suggest that higher confidence was more predictive of accurately detecting high-pleasant target stimuli on the forearm versus the palm, indicating higher metacognitive sensitivity, i.e., affective touch awareness, on CT than on non-CT skin (albeit, only at a trend level in our second experiment). This finding seems compatible with the social touch hypothesis suggesting that the CT system may be specialised for affective touch awareness, allowing individuals not only to discriminate between affectively relevant and irrelevant information (level 1 performance) but also to be aware of the accuracy of such discriminations (particularly in adults that presumably have a lifetime of experience with affective touch) in order to be able to guide action. This interpretation would imply that some domain-specific processes contribute to affective touch awareness, perhaps by rendering affective touch stimuli particularly salient, which is not compatible with the supramodality of metacognition, an ongoing debate in the field. However, before drawing any firm conclusions on this issue, further studies should compare metacognition of affective touch stimuli against other modalities in the same population and use larger datasets that will allow the use of other measures such as meta d’ (Fleming & Lau, 2014). Future studies should examine other drivers of confidence in participant’s task performance (Denison, 2017), as well how individuals may ‘learn’ to discriminate between stimuli over-time.

To the best of our knowledge, this is the first study to combine affective touch with signal detection paradigms and hence confirm the alleged role of the CT system in discriminating between affective relevant and non-relevant information. However, the present findings should be considered in light of certain methodological limitations. First, our SDT design employed two predetermined thresholds, whereas SDT research typically involves only one predefined threshold. For example, in experimentally induced pain, participants are asked to identify pinprick pain as ‘high’ or ‘low’ based on a predetermined pain threshold (i.e., the moment participant’s begin to feel sharp pinprick pain, e.g., Borhani, Làdavas, Fotopoulou, & Haggard, 2017). In contrast, our SDT design employed two thresholds: below and above the CT range. This was done because of two main reasons. First, in pain for example, an increase in noxious stimulation leads to a linear increase in pain perception, while an increase in tactile velocity does not lead to an increase in perceived pleasantness, but rather to an increase in tactile acuity. Second, this was done in order to ensure that participant’s pleasantness discrimination was not merely based on the increasing velocity of the touch. In fact, if this had been the case, we would have observed d’ values very close to 0. Thus, although the two noise distributions (i.e., touch velocities below and above the CT ranged) were grouped together in order to analyse the data according to the SDT principles (with the same number of trials for noise and signal), future research should examine whether these results hold when setting only one threshold. For example, one could predetermine the threshold at 1 cm/s and examine the discrimination between speeds within the CT range (e.g., at 3 cm/s and 6 cm/s) and below the CT range (e.g., .3 cm/s and .5 cm/s), or the other way around. This was not further explored in the current data given the low number of trials (i.e., doing this would result in only 8 noise and 8 signal trials per skin area), necessitated by the repetitive nature of the task and the fatigue properties of the tactile system. Indeed, the number of trials for the SDT task is in the low side for measuring sensitivity d’ and response bias C, and even more for the assessment of metacognition (Green & Swets, 1966).

Second, a trained experimenter always delivered the touch in this study, but this increased the social relevance of all stimuli and could have led to some human error, particularly given the eight different velocities administered in these two experiments. In order to optimise precision, future research may want to deliver the touch by means of a robotic device, as in some previous studies using rating scales (Triscoli et al., 2013).

Finally, the experimenter providing the touch was always female, whereas the touch receivers were both females and males. In contrast to previous research suggesting gender effects associated with the perception of touch (e.g., Bendas et al., 2017; Ilona Croy et al., 2014; Essick et al., 2010; Gazzola et al., 2012) there were no gender effects found across any of our outcome measures, including our manipulation checks on pleasantness ratings (see Supplementary Materials). Future research is needed to investigate this issue in a full factorial design, i.e., manipulating the gender of the touch receiver as well as the gender of the touch provider.

In sum, our findings suggest that there is not only increased affective touch sensitivity on the forearm versus the palm (as reflected by sensitivity d’; type 1 task), but also, higher confidence judgments regarding subject’s performance on this task seem to better predict the accuracy on detecting the high-pleasant target stimuli on the forearm versus the palm (as reflected by the aROC analyses; type 2 task). The present study provides novel and strong psychometric support for the social touch hypothesis on affective touch by showing that people are better at discriminating affective from non-affective tactile stimuli in on a skin site that contains CT afferents than on a skin site that does not. Moreover, the current study suggests that people have a better subjective awareness regarding their affective discrimination ability on CT innervated versus non-CT skin, suggesting a novel, domain-specific, metacognitive hypothesis that can be further explored in future studies.

## Author Contributions

M. Von Mohr, L. P. Kirsch and A. Fotopoulou developed the hypothesis and research plan. J. Loh, M. Von Mohr, and L. P. Kirsch collected the data. M. Von Mohr and L.P. Kirsch analysed the data. M. Von Mohr and L. P. Kirsch wrote the manuscript, under the guidance of A. Fotopoulou. All authors approved the final version of the manuscript for submission.

## Acknowledgements

This work was supported by the European Research Council (ERC) Starting Grant ERC-2012-STG GA313755 (to A.F.) and the National Council on Science and Technology (CONACyT-538843) scholarship (to M.V.M.).

