## Supplementary Materials for "Affective Touch Dimensions: From Sensitivity to Metacognition"

### Supplementary Material

#### Experiment 1

*Gender effects*

We conducted a mixed 2 (skin area: forearm, palm) x 2 (gender: female, male) ANOVAs to examine whether there were any gender differences separately in our main outcome measures, i.e., sensitivity d', response bias C and aROC. Results suggest that there was no between-group main effect of gender, p's>.274 (except for a trend, p=.097 on sensitivity d'), and gender did not interact with skin area, p's>.319, in any of our outcome measures. Similarly, a 2 (skin area: forearm, palm) x 2 (speed: CT-optimal speeds; CT non-optimal speeds) x 2 (gender: female, male) ANOVA was also conducted on our manipulation check of pleasantness ratings. As above, there was no main effect of gender, p=.284, and gender did not interact with skin area, p=.672, speed, p= .363, or their combination, p=.674. Thus, these results suggest that there are no effects of gender associated with our main SDT task or our manipulation checks consisting of pleasantness ratings.

*Interim analyses*

Interim analyses where conducted on the first half of the sample (i.e., 47 participants) from Experiment 1. These interim analyses suggest significantly higher sensitivity (d') on the forearm (*M*=1.62, *SD*=.92), as compared to the palm (*M*=1.19, *SD*=.94), *t*(45)=3.16, *p*=.003, *d*=0.47, with a trend towards significance for the response bias, indicating more negative response bias (C) in the forearm (*M*=-.28, *SD*=.48), as compared to the palm (*M*=-.11, *SD*=.63), *t*(45)=-1.79, *p*=.081, *d*=-0.26. These are the same pattern of results shown in the main results.

*Manipulation check (pleasantness ratings)*

We conducted a 2 (skin area: forearm, palm) x 2 (CT-optimal speeds; CT non-optimal speeds) ANOVA on pleasantness ratings to examine whether on average participants experienced a CT-optimal velocity as more pleasant than a non-CT optimal velocity. Follow-up analysis used paired t-tests with Bonferroni correction when applicable.

Analyses conducted on the pleasantness ratings scores suggested that touch at CT-optimal speeds, versus non-CT optimal speeds, was perceived as expected at the group level. Specifically, across skin areas, participants perceived touch at CT optimal speed as more pleasant than touch at non-CT optimal speed, $F(1,93)=91.77$, $p<.001$, $\eta^2_{partial}=.49$. Similarly, participants perceived the touch as more pleasant on the forearm, versus the palm, $F(1,93)=5.90$, $p=.017$, $\eta^2_{partial}=.06$. Importantly, skin area interacted with speed, $F(1,93)=4.45$, $p=.038$, $\eta^2_{partial}=.05$. Post-hoc tests, using Bonferroni adjusted alpha level of .0125 per test (.05/4), showed that participants perceived touch at CT-optimal speed (forearm: M=69.73, SD=14.98; palm: M=64.95, SD=14.77) as more pleasant than touch at non-CT optimal speeds (forearm: M=46.23, SD=18.34; palm: M=45.03, SD=19.58) on both forearm, $t(93)=10.31$, $p<.001$, and palm, $t(93)=7.80$, $p<.001$, respectively. Importantly, while there was no difference between palm and forearm in the perceived pleasantness of touch at non-CT optimal speeds, $t(93)=.79$, $p=.431$, participants perceived touch at CT optimal speeds as more pleasant in the forearm as compared to the palm, $t(93)=3.22$, $p=.002$.

**Experiment 2**

*Gender effects*

We conducted a mixed 2 (skin area: forearm, palm) x 2 (gender: female, male) ANOVAs to examine whether there were any gender differences separately in our main outcome measures, i.e., sensitivity d', response bias C and aROC. Results suggest that there was no between-group main effect of gender, p's>.227, and gender did not interact with skin area, p's>.145, in any of our outcome measures. Similarly, a 2 (skin area: forearm, palm) x 2 (speed: CT-optimal speeds; CT non-optimal speeds) x 2 (gender: female, male) ANOVA was also conducted on our manipulation check of pleasantness ratings. As above, there was no main effect of gender, p=.750, and gender did not interact with skin area, p=.687, speed, p= .319, or their combination, p=.946. Thus, similar to Experiment 1, these results suggest that there are no effects of gender associated with our main SDT task or our manipulation checks consisting of pleasantness ratings.

*Manipulation check (pleasantness ratings)*

We conducted a 2 (skin area: forearm, palm) x 2 (CT-optimal speeds; CT non-optimal speeds) ANOVA on pleasantness ratings to examine whether on average participants experienced a CT-optimal velocity as more pleasant than a non-CT optimal velocity. Follow-up analysis used paired t-tests with Bonferroni correction when applicable.

Analyses conducted on the pleasantness ratings scores suggested that touch at CT-optimal speeds, versus non-CT optimal speeds, was perceived as expected at the group level. Specifically, across skin areas, participants perceived touch at CT optimal speed (M= 65.10, SD=11.62) as more pleasant than touch at non-CT optimal speeds, (M=44.91, SD= 12.86),

$F$(1,98)=202.65, $p$<.001, $\eta^2_{partial}$=.67. There was a trend for a main effect of skin area, $F$(1,98)=3.68, $p$=.058, $\eta^2_{partial}$=.04, and skin by speed interaction, $F$(1,98)=3.64, $p$=.059, $\eta^2_{partial}$=.04.
